# The Giant Clam Photosymbiosis is a Physically Optimal Photoconversion System for the Most Intense Sunlight on Earth

**DOI:** 10.1101/2023.02.28.530416

**Authors:** Amanda L. Holt, Lincoln F. Rehm, Alison M. Sweeney

## Abstract

Giant clams are photosymbiotic with unicellular algae (“zooxanthellae”) organized in the clam’s mantle tissue. This tissue has an especially low albedo for a photosynthetic system, generally less than 10% at all visible wavelengths. This efficient absorbance of light occurs in the ecological context of the high solar irradiances in intertidal habitats near the equator. At these light levels, photosynthetic systems typically adapt to absorb less light in order to prevent radiative damage to chloroplasts. Giant clams are therefore unusual. If the giant clam photosymbiosis proves to be simultaneously efficient at absorption and at phototransduction at these irradiances, they are potentially remarkably productive and an important source of bioinspiration. We showed previously that the clams organize algae into vertical pillars in the mantle tissue. The clams’ iridocytes, or optically structured skin cells on the surface of the tissue, then function to evenly distribute incoming solar irradiance along the vertical faces of the pillars. The result is that zooxanthellae in the system absorb solar power at lower rates than that of incoming solar flux. The overall energetic performance of this phtooconversion scheme has, however, been difficult to characterize given the complex three-dimensional structure and the fact that it is coupled to a much more voluminous, respiring animal. Here we use a combination of photochemical characterization and new quantitative modeling of data from the literature to estimate the photochemical efficiency as a function of incoming irradiance of the initial electron-transfer events.

Our approach is to consider the clam mantle tissue in isolation as a meta-material for photoconversion. To do this, we developed a method to directly measure the system’s photochemical efficiency with spatial resolution of 10’s of microns using optical microprobes threaded through the tissue. These experimental efficiency data then serve as ground-truthing for a subsequent reanalysis of photosynthesis-irradiance curves of clams taken from the literature. For this quantitative re-analysis, we incorporated the clam system’s quantum efficiency as a function of irradiance per cell into a Monte Carlo model of radiative transfer among cells to find the tissue’s area-specific oxygen evolution apart from any sinks. We found that cells located within the dense clam system had fluorescence transients (i.e., Kautsky curves ^1^), a direct measure of the efficiency of PS II) that were very slow and of low intensity, particularly for a dense system, consistent with photochemical efficiencies generally greater than 50% and often greater than 90%. When incorporated into a larger computational model, we found that mature Tridacnid clams can efficiently perform photoconversion of light energy into chemical energy at light intensities many times more intense than the maximum time-averaged environmental radiance, or even the solar constant. The intensities to which the clam is adapted, however, can be found in strong wave-lensed pulses of irradiance that are characteristic of the clams’ habitats. This surprising result makes sense if the system has evolved to both avoid damage from and utilize the power in the intense pulses of light that result from wave-lensing. Our model predicts that by evolving to compensate for the intense pulses of solar energy produced by wave-lensing, the clam system can perform photochemical conversion of radiation at intensities many times greater than the solar constant at around 90% quantum efficiency. This result in turn suggest a strategy for engineered organic and biological composite materials performing photoconversion under solar concentration.

**Graphical Abstract:** 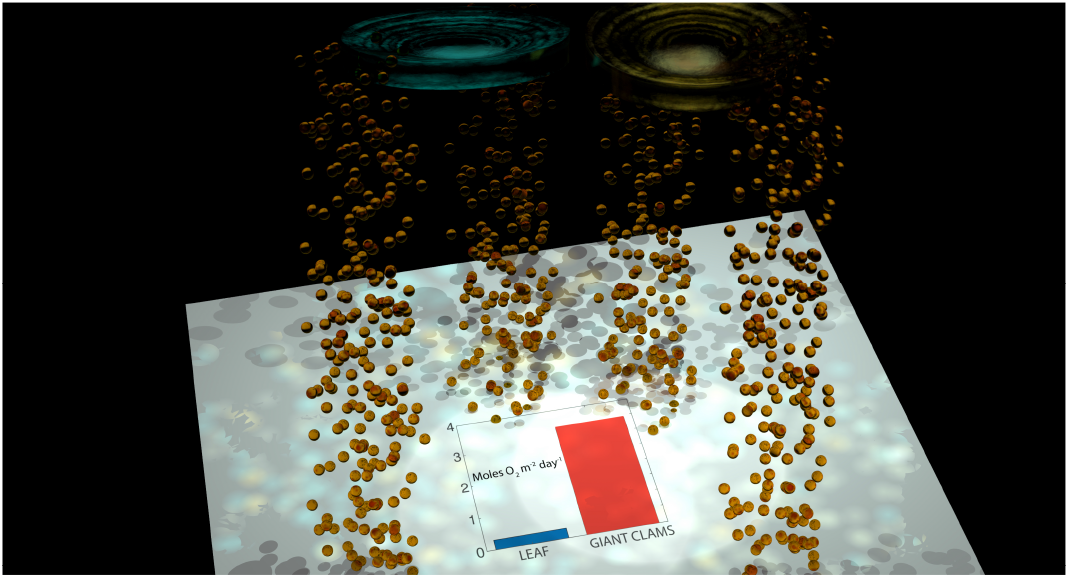

## Introduction

We have previously characterized the structure of the photosymbiotic mantle tissue of giant clams living on shallow, clear Indo-Pacific reefs^2^ (Figure 1,2). We showed that the symbiotic dinoflagellates, or zooxanthellae, responsible for the system’s photosynthesis are organized into pillars roughly 100 μm in diameter, 100 μm apart, and a few millimeters long. In turn, the surface of the tissue is coated with iridocytes, or optically structured clam skin cells. The iridocytes act to evenly scatter the solar irradiance from the surface of the clam to the underlying pillars of algae.^2^ The resulting photon flux at the surface of the vertical algal pillars is then about 10% of the incident surface irradiance. In this way, all of the radiant power incident on the surface of the clam is absorbed by zooxanthellae at a much lower power. In principle, this downregulated power per cell could allow the symbiosis to perform photoconversion of solar energy near unitary quantum efficiency, as photosynthesis can operate extremely efficiently at low incident power.^3^ The time-averaged downwelling irradiance on the shallow, Indo-Pacific reefs where clams are found is around 1000 μE m^−2^s^−1^at noon. In addition, this environment experiences strong wave-lensing, or focusing of downwelling light through small surface waves under breezy conditions, resulting in intense flashes of irradiance (Figure 1). In the top few meters of the water column, these focused flashes can last 10’s of milliseconds with intensities ten times greater than the average background radiance.^**?**^

**Figure 1:**
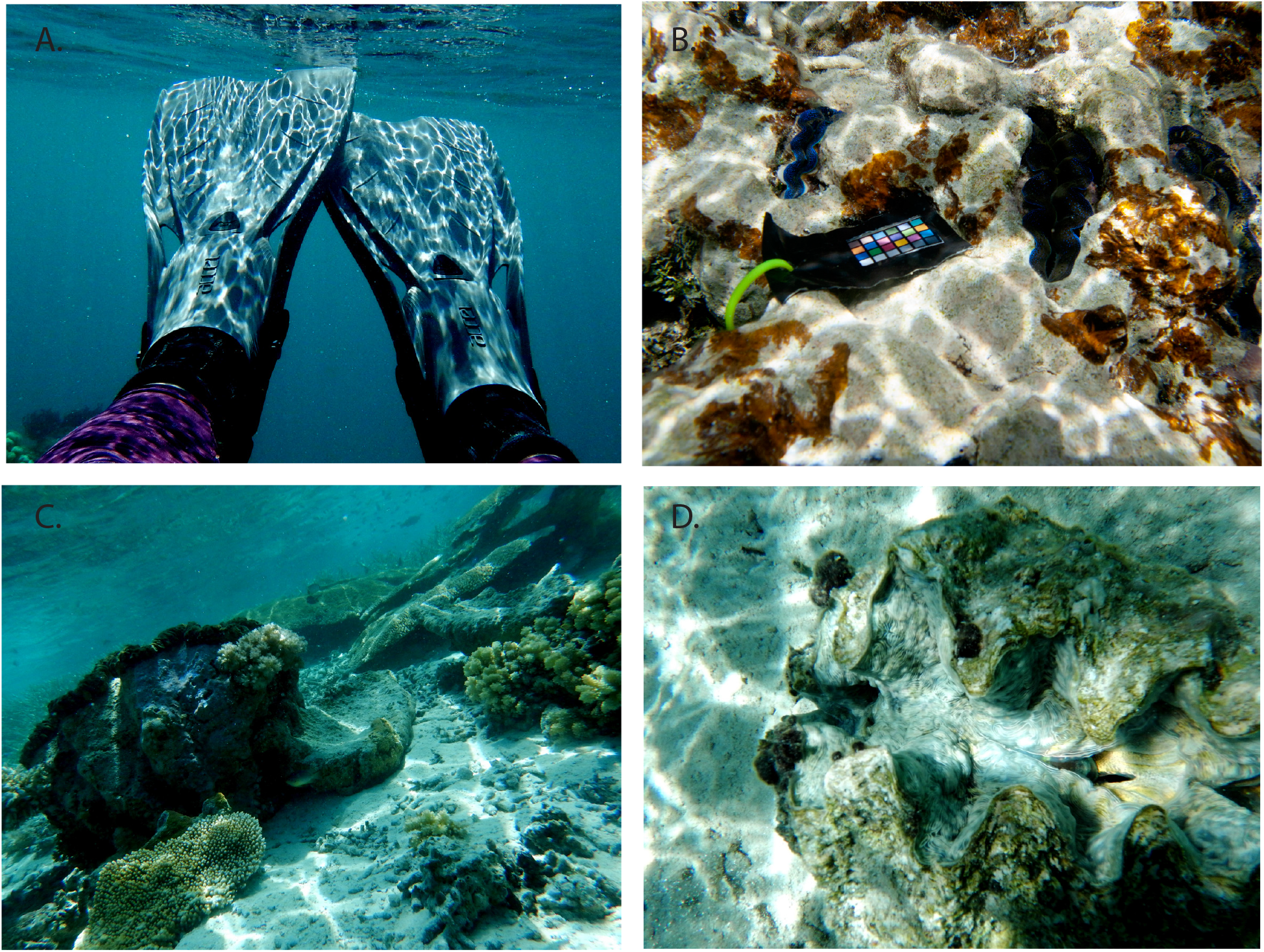
Giant clams in their strongly wave-lensed environment. (**A**) Wave-lensing in the first few centimeters of the water column as visualized on dive fins. The patterns due to lensing are extremely intense, even when reflected from a black background, and spatiotemporally frequent. (**B**) *Tridacna crocea* growing in the top of a coral head with wave-lensing. (**C**) *Tridacna gigas* on a shallow reef, notice the summation of wave-lensing regions on the sand near the animal. (**D**) *Hippopus hippopus* with camera-saturating, summing wave-lensed pulses on the nearby sand.

**Figure 2:**
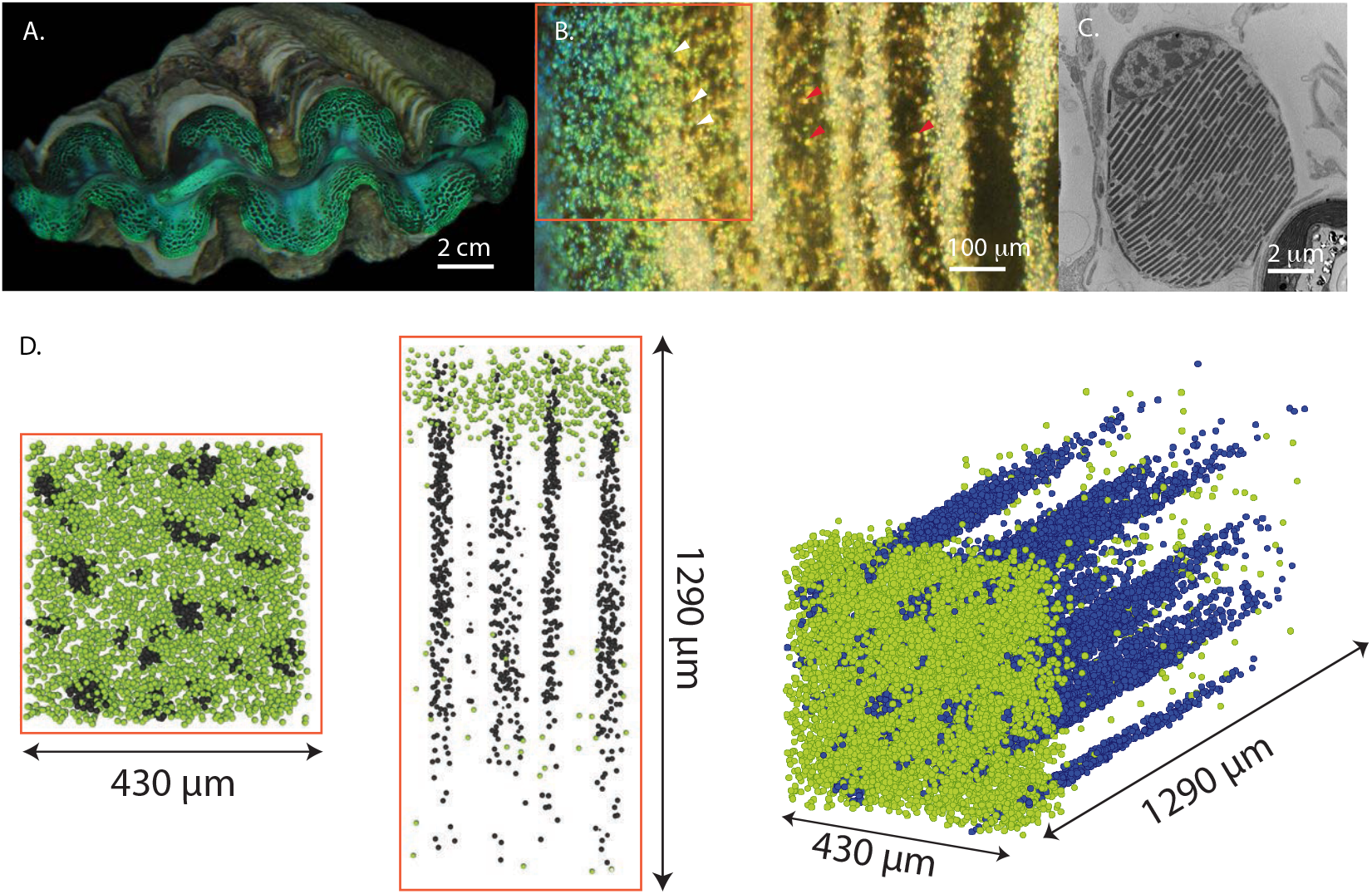
Overview of the tridacnid ‘giant’ clams and the physical model of their tissue.(**A**) Photograph of *Tridacna crocea* with green iridocytes. (**B**) Darkfield micrograph showing example iridocytes cells (red arrowheads) and tips of algae pillars emerging from underneath the layer of iridocytes (white arrowheads) (**C**) Single clam iridocyte in TEM (**D**) Modeled tissue sections; top view showing algae (grey dots) and iridocytes (green dots), side view (**F**) and three-dimensional view showing full scope and dimensions of modeled tissue

Previous investigators measured photosynthesis-irradiance relationships for living clams, and uncovered some paradoxes and uncertainties around the system’s photosynthetic performance.^4–12^ Past work on the system was motivated by the physiological symbiotic relationships between the host clam and the algal symbionts, and the resulting analysis accordingly considered respiration and total wet weight. This approach had the effect of obscuring the physical performance of the isolated mantle tissue regions of the organism as a material evolved for phototransduction. On closer examination, several interesting observations come to light. First, for large clams, oxygen evolution from symbionts never became saturated, even at irradiances of 3000 μE m^−2^s^−1^, or a light intensity three-fold greater than the time-averaged noontime radiance.^4^ The scaling between the density of algae in the clam’s mantle layer and the volume of the clam is also unusual–the number of algae in the system increases with an exponent of 0.99 of the clam’s wet weight, not 0.66 as would be the case if algae simply increased proportional to mantle surface area.^4^ This is an indication that the micropillared structures containing zooxanthellae that we described grow deeper into the tissue as a clam’s shell length increases. Experimental measurements of photosynthesis-irradiance curves of zooxanthellae, when extracted and isolated from the host, were both unpredictable and had high variance, were generally close to previous observations for planktonic algae, demonstrating saturated photosynthesis-irradiance curves at the usual light intensities. This result suggests both high underlying physiologic variability of zooxanthellae when hosted by the clam, and the presence of variable radiance within the host. We have also observed that healthy clams shed copious fecal pellets composed almost entirely of intact, viable algae. Therefore, the system seems to have excess, “invisible” structure and productivity, and growth of the clam itself is potentially a poor proxy for the overall productivity of the symbiosis.

Similarly, standard measurements of quantal photosynthetic efficiency, such as pulse-amplitude modulated fluorometry (PAM), provide stimulating illumination of the surface of a photosynthetic system. These techniques implicitly assume a dilute or two-dimensional geometry of absorberes in a system, and that there will be minimal influence of self-shading, multiple scattering, or re-absorption within the system. Our prior modeling and measurements of radiative transfer within the clam^2^ show a zone of relatively high irradiance within the top 100 μm of the tissue, followed by a decay in irradiance to an asymptote at an intensity about 10% of that incident on the surface of the tissue. It is this irradiance that is experienced by the majority of the cells in the system. So, in a densely structured system like the clam, the signal from an externally-stimulating experiment like PAM will necessarily be dominated by the most irradiated, most fluorescent zooxanthellae at the surface, which in this case represent a small fraction of the larger system and a small fraction of the total power flowing through the system. To understand photosynthetic efficiencies inside the clam symbiosis, we adapted our previous microprobe technique for measuring internal tissue irradiance^2^ to measure fluorescence transients as a direct output of photosynthetic quantum efficiency within the living clam tissue.

Next, we used this information to develop newly detailed models of the clam system’s productivity in different irradiance contexts. First, we re-utilized the whole-clam photosynthesis-irradiance curves from the work by Fisher and colleagues to find an average per-cell quantum efficiency as a function of irradiance of the clam system.^4,13^ We calculated the system’s integrated productivity over a circadian cycle of photosynthetically active radiation (PAR) averaged over short time scales (seconds) but variable over the day. We also estimated the system’s instantaneous performance as a function of the intense pulses of irradiance caused by wave-lensing, and found the intensity at which this model predicts photoinhibition.

Our results show that the giant clam system has evolved to perform photoconversion at near-optimal quantum efficiency while absorbing irradiance at intensities at least ten-fold greater than noontime downwelling irradiance in the tropics, i.e., conditions of solar concentration. Clams experience pulses of these extreme intensities several times a minute due to wave-lensed caustics in shallow, clear tropical water, providing a force for evolutionary adaptation. There is a close match between the characteristics of wave-lensed flashes in the ocean and the structure and photosynthetic performance of the clam. Clams are, therefore, an important system for learning the fundamentals of using organic materials for efficient solar photoconversion under solar concentration such as biofuel production and organic photovoltaics.

## Methods

### Double optical-microprobe construction

To measure photosynthetic transients within the highly absorbing living tissue, we modified an optical microprobe technique to incorporate two optical fibers. One of these fibers stimulates the tissue with 455 nm light, and the second fiber receives the resulting time-resolved chlorophyll fluorescence response (Figure 3).

**Figure 3:**
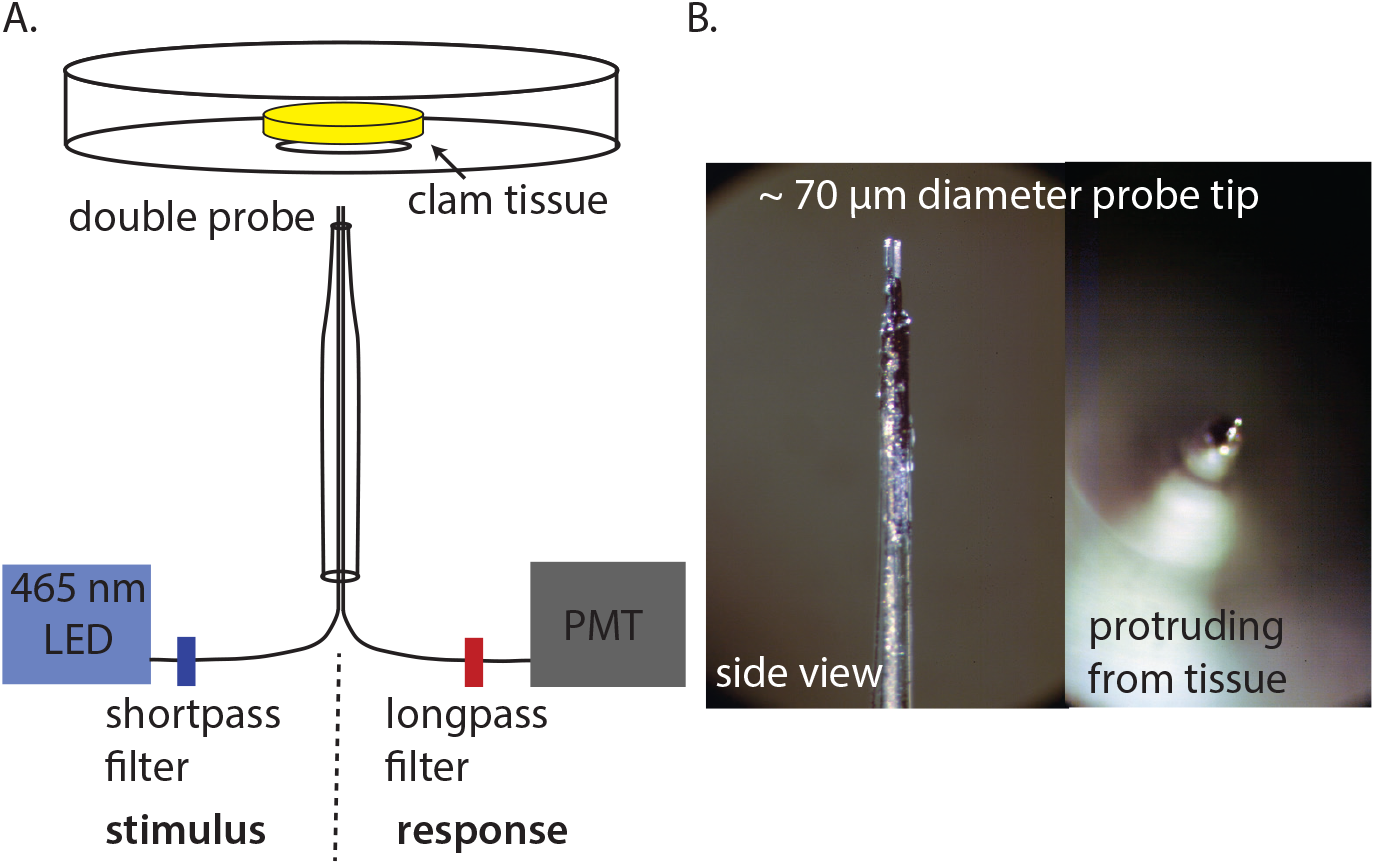
Experimental setup for measuring fluorescence transients within strongly absorbing, living clam tissue. (**A**) Diagram depicting the double microprobe and sampling geometry. One of the two pulled optical fibers serves the stimulus half of the experiment, delivering light from a 455 nm LED through a short-pass filter and into the clam tissue through a fiber optic with a tip diameter of about 35 μm. The response fiber conducts long-wave light from chlorophyll fluorescence from the tissue, through a long-pass filter, and into a photomultiplier tube coupled to an oscilloscope (**B**) Photographs of the side and top view of the microprobe.

### Tissue preparation and fluorescence transient measurement

We measured fluorescence transients within the living tissue of several species of giant clam. Tridacna derasa, T. maxima, T. crocea, Hippopus hippopus, T. squamosa and T. gigas were purchased from clam aquaculture facilities in Palau and kept alive in running seawater at the Palau International Coral Reef Center (PICRC) in Koror, Palau. 8-mm diameter biopsies of clam mantle tissue were immobilized in a petri dish using gelatin.

The Petri dish containing the clam sample was then mounted on a stage attached to a micromanipulator and positioned to center the hole in the bottom of the dish around the fiber optic probe tip, which was mounted vertically and held stationary throughout the experiment. We then lowered the dish in fixed increments onto the double-optical probe, pushing the probe through the clam tissue (Figure 3A).

Stimulating 455 nm light was provided by a fiber optic-coupled high power LED coupled to one of the two fibers in the probe. The fluorescence transient was measured in turn using the second fiber from the probe connected to the SMA port of a photomultiplier tube. The photomultiplier tube (PMT) signal was detected using an oscilloscope.

We measured fluorescence transients at different light stimulation intensities of 2000, 3500, and 5000 μE m^−2^s^−1^at the immediate surface of the probe. These stimulating intensities are higher than typically used in this type of experiment, however, in practice these intensities were required to generate measurable fluorescence responses specifically when measuring zooxanthellae within clams. For terminological convenience, we considered 2000 μE m^−2^s^−1^at the probe tip to be “actinic” and 5000 μE m^−2^s^−1^at the probe tip to be “saturating” in our subsequent analysis of these data, while also noting that these terms are typically defined at much lower intensities. Prior to each measurement, the clam tissue sample was dark adapted at room temperature under a box for 15 minutes. Due to the sensitivity of the PMT and to ensure all signal was from chlorophyll fluorescence, all measurements were performed in a dark room.

### Fluorescence transient data analysis

To calculate the maximum quantum yield of zooxanthellae *in hospite* resulting from these measurements of the fluorescence transient, we started by identifying the zero-time point of each measurement where signal started to rise by eye. Then, using purpose-written Matlab code, we used a median filter to smooth the rapidly sampled oscilloscope data while preserving the time-dependence of the fluorescence signal. The quantum efficiency of the algae near the optical probe is characterized by the minimum chlorophyll fluorescence at the zero-time point (*F*_0_) and the maximum fluorescence at a time greater than 1 s (*F_M_*). To find *F*_0_ from the fluorescence transient data, we made a linear fit to the fluorescence signal between 40 and 120 μs and calculated its intercept at the zero-time point.^14^ *F_M_* is then the steady-state fluorescence value after 1 s when the signal was averaged over 5 ms intervals. The maximum quantum yield, *q* is then calculated using the definition 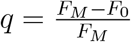^1^ (Figure 4).

**Figure 4:**
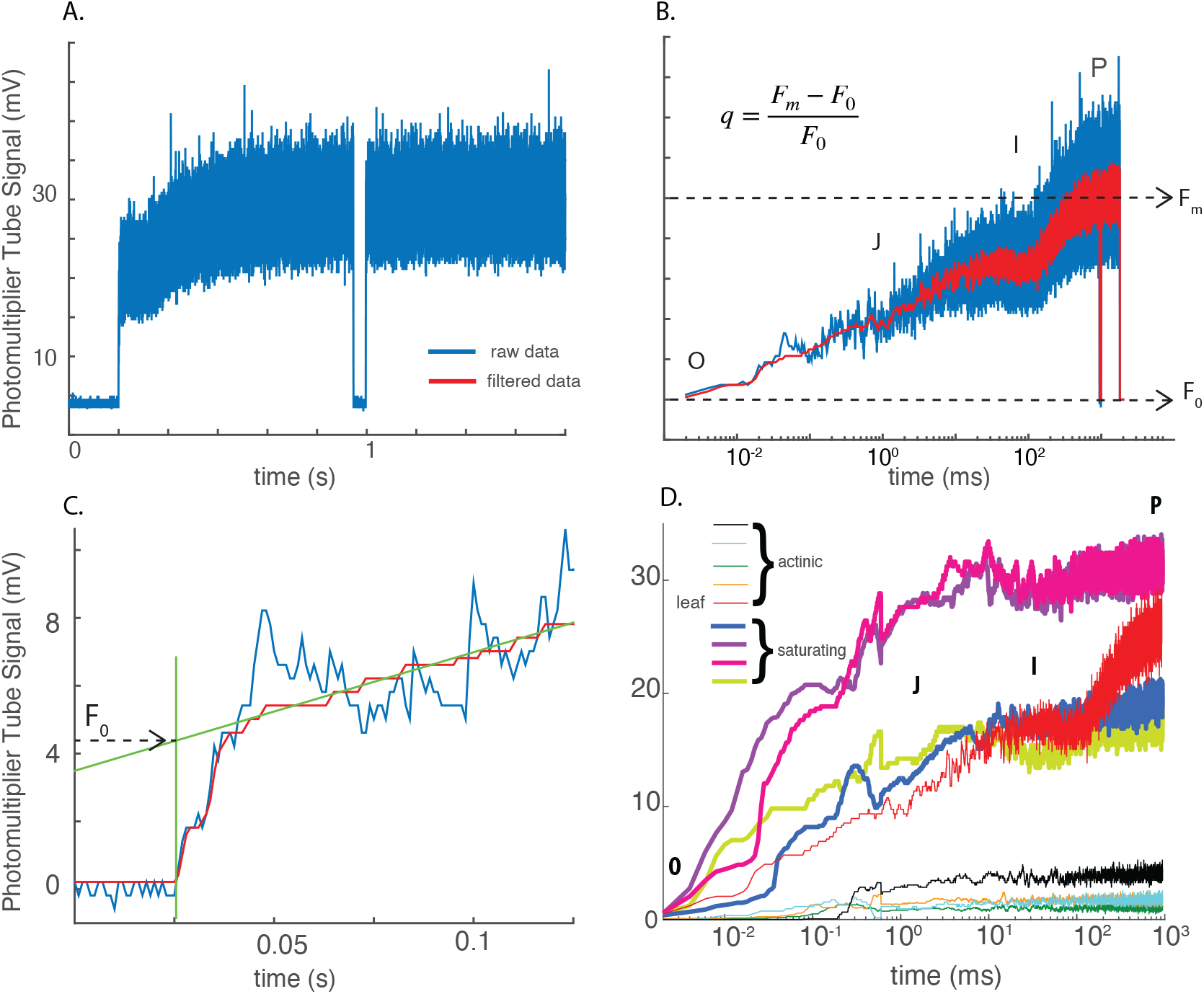
Fluorescence transient methods and sample data. (**A**) Example raw transient curve (**B**) Raw transient data in log time (blue curve) and after smoothing (red curve) and example calculation. The positions OJIP are labeled and the calculation for the maximum quantum efficiency is presented with positions of the maximum fluorescence (F_m_) and the minimum fluorescence at time=0 (F_0_) (**C**) Example curve on technique used for finding F_0_. (**D**) Example *in situ* fluorescence transient curves from clam tissue. Bold curves are sample data taken at saturating light levels while thin curves are taken at actinic light levels.

### Monte-Carlo Modeling

The Monte Carlo algorithm used in this paper is described in detail by Holt et al.^2^ Briefly, we generated three-dimensional coordinates of algae and iridocytes based on two-dimensional panoramic light micrographs and dark field light microscope images. These coordinates were used in a Monte Carlo algorithm of light propagation where each cell type scattered photon packets according to modeled phase functions based either on Mie scattering (algae) or Discrete Dipole Analysis (iridocytes).^2,15^ Using this Monte Carlo model, we calculated the average per-cell absorption as a function of tissue depth for three scenarios, keeping all geometric parameters the same except for the maximum depth of the tissue. We generated three-dimensional cell coordinates for algal pillars of about 4.6 mm depth, consistent with large individual clams (1000’s of grams wet weight). We then truncated these coordinates twice, once at 1.2 mm (medium-sized clam with 18.2 g wet weight) and again at 0.350 mm (small-sized clam with a 0.014 g wet weight), such that the positions of algae and iridocyte cells were equivalent for each model up until the maximum tissue depth. This set of models is consistent with our observations of clams of different size- and age-classes, where the two-dimensional arrangement of the pillars within the mantle tissue doesn’t change as clams grow, but the mantle tissue becomes thicker with shell length. It is also consistent with the scaling relations between shell length, wet weight and zooxanthellae cell number cited in Fisher et al.^4^ The calculated whole-tissue abosorbance of these model systems is also in agreement with experimental measurements of whole-tissue absorbance measured by doubled integrating spheres, described below.

### Clam tissue and single-cell absorbance measurements

To experimentally measure the total amount of light absorbed in the clam mantle tissue, we used a doubled integrating sphere approach. We prepared mantle biopsies embedded in gelatin as described above and in the supporting information, and sandwiched the slab of gelatin containing the clam tissue between two high-precision windows with antireflection coating. This preparation was then pressure mounted between two integrating spheres. The light source for this experiment was a fiber-optic coupled white LED secured to one integrating sphere. Light measured from the sphere with the light source mounted is then primarily back-scattered from the sample, and light measured from the sphere without the light source is light primarily transmitted by the sample. Fibers were mounted perpendicular to the light path on both integrating spheres to characterize this back-scattered and forward-scattered light. We standardized this measurement of clam tissue by characterizing the light in both spheres without a clam sample present. The signal missing in the second sphere when the clam tissue is present, relative to the signal when it is absent, is then the light that is absorbed by the tissue. We also characterized the absorptance of several green leaves in this manner (Figure 5A,B).

**Figure 5:**
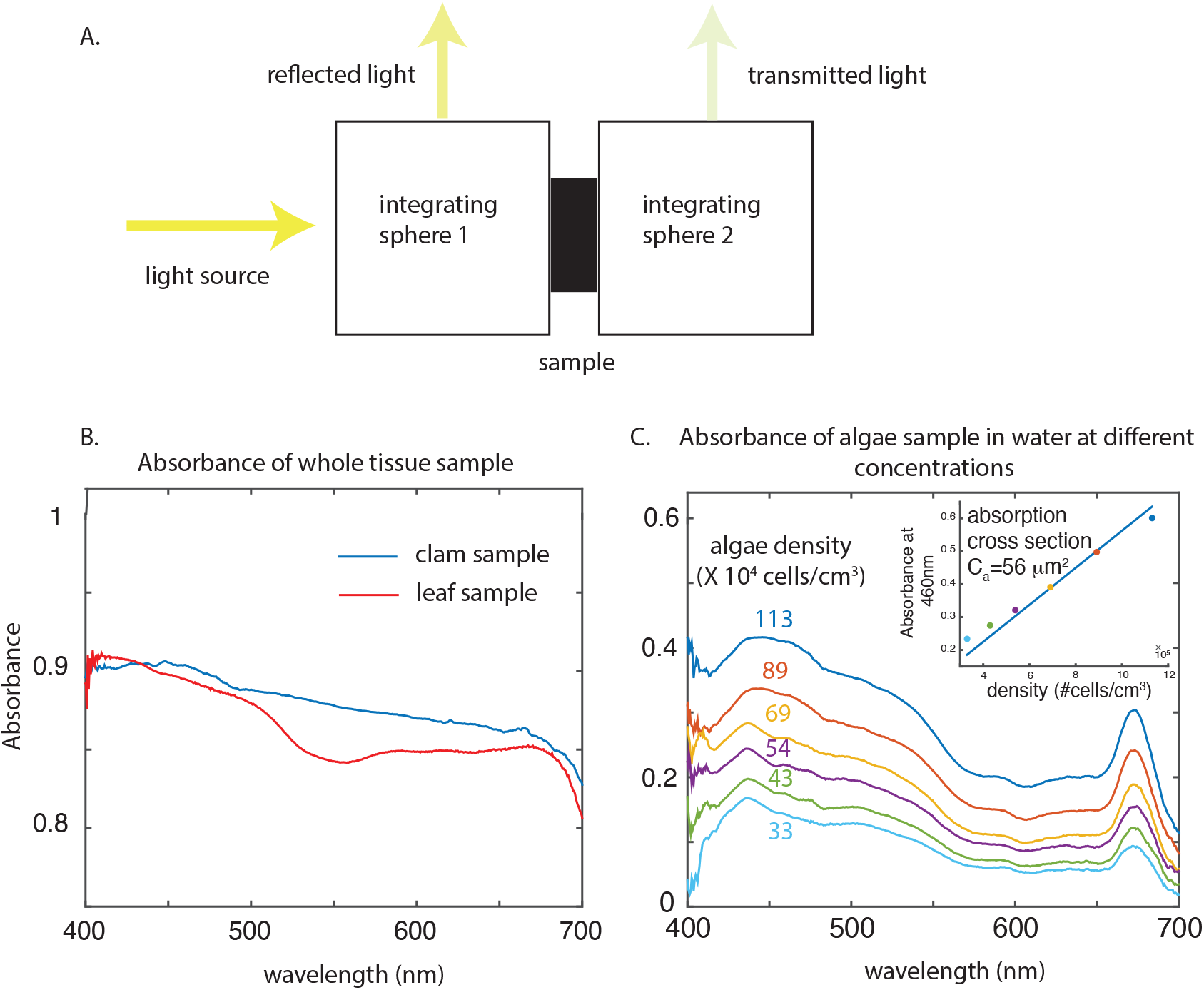
Double integrating sphere experimental method and absorption data from clam tissue and algae in a cuvette (**A**) Schematic of double integrating sphere setup. (**B**) Representative measurements of the absorbance of clam tissue and a green leaf obtained by the double integrating sphere apparatus. (**C**) Absorbance data for increasing concentrations of algae suspended in solution. Insert shows calculation of absorption cross section of algae cells that is subsequently used in productivity model.

We also estimated the absorptance and transmittance of single zooxanthellae cells from a dilute suspension of cells. Cell suspensions were then placed in a 1 cm-pathlength quartz cuvette mounted between two integrating spheres (Figure 5A). The total reflectance and transmittance of the algae in solution was measured from the signal in the two spheres and normalized to a quartz cuvette filled with clean growth medium. The cell number density of each dilution was measured using a hemocytometer. We calculated the single-cell absorption cross-section from the least-squares solution to the absorbance (log of transmittance values at 460 nm) versus concentration data points (Figure 5C).

### Primary productivity model

We sought to understand how the clam’s system of spatially structured scatterers and absorbers changes the photosynthetic yield of the system relative to that of a spatially homogeneous system of the same average cell density. In general, the oxygen evolution rate (OER) for a photosynthetic system is estimated to be *PAR* × *a* × *Q* × 0.5 × 0.25 where PAR is photosynthetically active radiation incident on the system, *a* is the percent of PAR absorbed by the system, *Q* is the photosynthetic quantum yield, the factor of 0.5 is an estimate of the fraction of absorbed PAR going to photosystem II, and the factor of 0.25 is estimated to be the number of O_2_ molecules evolved per electron given four electrons for one O_2_ molecule.^16^

We expanded this method of estimating OER to account for variable light intensities, wavelengths, and single-cell absorbances as a function of tissue depth and time of day:

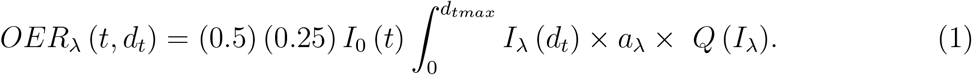

Here, *d_t_* is the depth below the surface of the tissue; *I*_0_ (*t*) is the incident PAR on the surface of the clam at time *t*; *I_λ_* (*d_t_*) is the intensity of light of wavelength λ present at tissue depth *d_t_* up to *d_tmax_*, the maximum modeled depth; *a_λ_* is the percentage of radiation absorbed by a single cell at wavelength λ; and *Q* (*I_λ_*) is the quantum efficiency.

In this calculation, we extrapolated cellular quantum efficiency (*Q*(*I*λ)) from photosynthesis-irradiance (PI) relationships of the clam system reported in the literature using the method of Eilers and Peeters.^4,13,17,18^ We then normalized the efficiency curves to maximum efficiency at zero irradiance.

We used this approach to finding quantum efficiency as a function of irradiance to calculate productivity from four different model tissues: 1) homogeneously distributed algae in a volume of 0.5 × 0.5 × 4.6 cm^3^ as in a pelagic system; 2) tissue consistent with a 0.14 g clam with a model volume of 0.5 × 0.5 × 0.35 cm^3^; 3) tissue consistent with a medium-sized clam of mass 18.2 g and a model volume 0.5 × 0.5 × 1.25 cm^3^; and 4) tissue consistent with a large-size clam of mass 1700 g and modeled volume 0.5 ×0.5 × 4.6 cm^3^) (Figure 7D, 6B). This is a relatively conservative approach since the experimental values of quantum efficiency we measured *in hospite* using photosynthetic fluorescence transients in the central regions of the tissue were generally much higher, and at higher irradiances in central regions. In contrast, this approach using PI curves from the literature incorporates respiration by the host and is a system average, including less efficient cells at high irradiances at the top of the tissue. Therefore, we anticipate that productivity estimates resulting from this model of quantum efficiency likely underestimates the true performance of the system.

**Figure 6:**
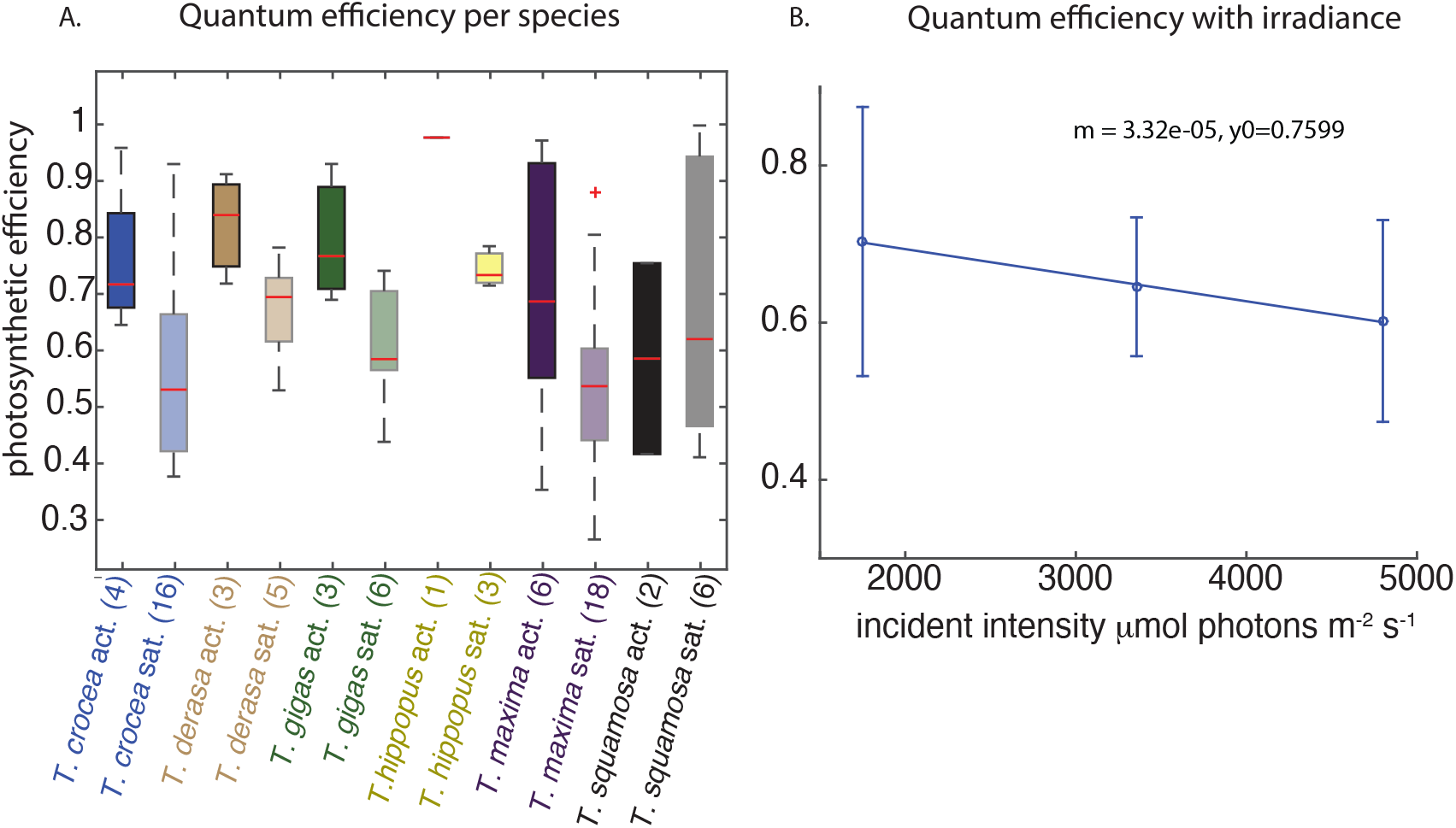
Within-tissue maximum quantum efficiency (**A**) Box plots showing averages and standard deviations of maximum quantum efficiency as measured using the double probe by species. Measurements under actinic irradiance (2000 μE μE m^−2^s^−1^) and saturating irradiance (5000 μE m^−2^s^−1^are shown. Values in parentheses indicate the number of measurements represented by the box plot. (**B**) Maximum quantum efficiency with incident light level with error bars and slope calculation.

**Figure 7:**
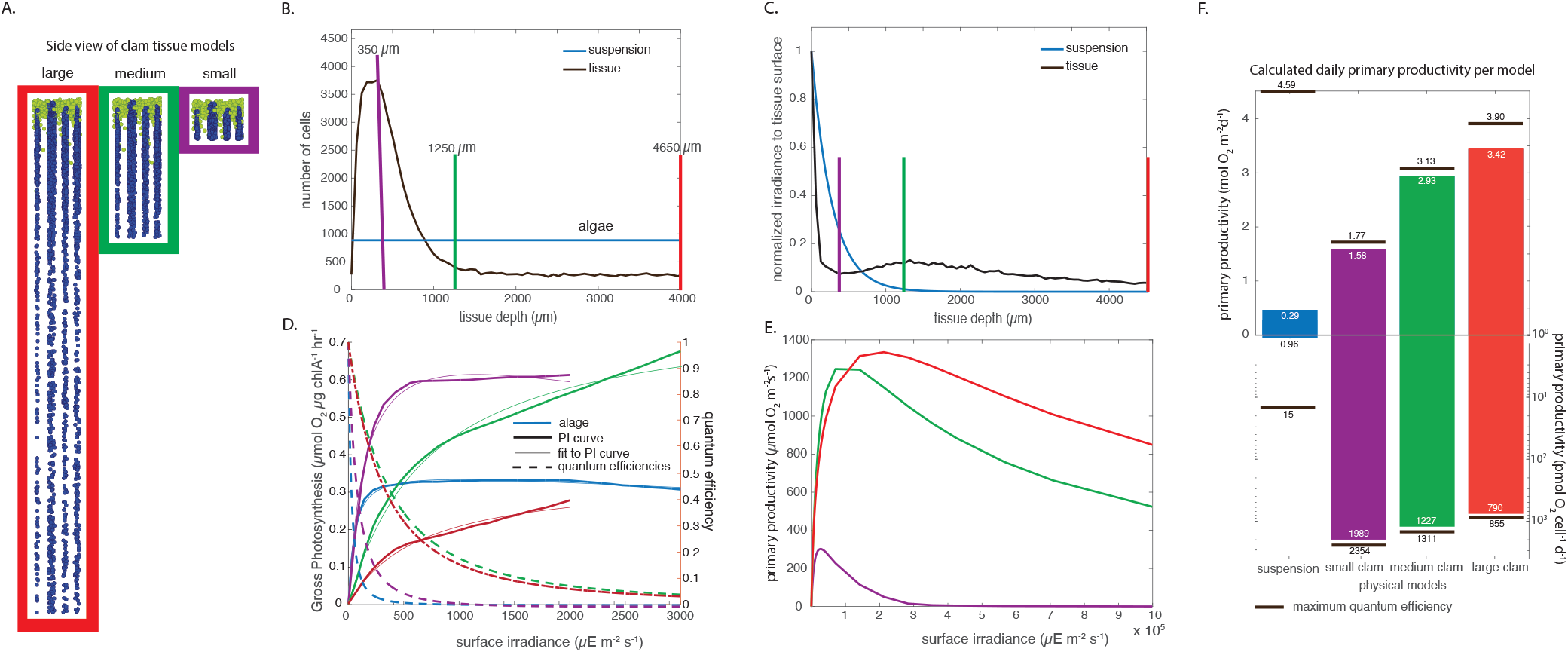
Productivity models: Inputs to the model include cell density, irradiance with tissue depth and photosynthetic efficiency with irradiance. (**A**) Side view of the cell positions of the three clam models used in the productivity calculation: small (0.5×0.5×0.35 mm^3^, purple), medium (0.5× 0.5× 1.25 mm^3^, green) and large (0.5× 0.5× 4.6 mm^3^, red). Iridocyte cell positions are shown in green, and zooxanthellal cell positions are shown in blue. (**B**) Algae cell density in the clam tissue models with tissue depth. Vertical lines indicate the bottom of the tissue for the small (purple), medium (green), and large (red) clam models. (**C**) Normalized irradiance with tissue depth. Vertical lines indicate the bottom of the tissue for the small (purple), medium (green), and large (red) clam models. (**D**) Gross photosynthesis with irradiance for different sized clams (PI curves, bold lines);^4^ corresponding quantum efficiencies (dashed lines), and the corresponding back-calculated fits to the PI curves (thin lines). (**E**) Calculated primary productivities with fixed surface irradiance values for three clam models up to irradiance of 1,000,000 μE m^−2^s^−1^. (**F**) Daily total productivity for each clam model. Top graph shows oxygen evolution normalized by area and graph shows data normalized per cell. Black lines and labels show the physical optimum if every cell has a quantum efficiency of 100%. White numbers show the value corresponding to the bar’s length for the clam’s performance.

For the clam system with spatially organized cells, *I_λ_* (*d_t_*) was determined using our Monte Carlo model of radiative transfer within the clam tissue. For a system of homogeneous average cell density, the irradiance as a function of tissue depth will be:

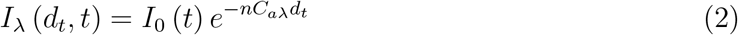

where *n* is the number density of algae cells, set to be equal to the average density in the clam tissue, *C_aλ_* is the absorption cross section of algae cells determined from measuring the absorbance of a dilute suspension of cells, and *d_t_* is the depth in the algae layer and is also set equal to that of the clam, and *t* is time.

We obtained the time-dependent environmental PAR (*I*_0_ (*t*)) from a database from the Pacific Islands Ocean Observing System.^19^ We calculated *OER*_*λ*(*t*)_ over a 24-hour period using *Q* (*I_λ_*) for the four systems mentioned above.

To model the effects of intense pulses from wavelensing and the performance of the system at intensities higher than short-time averaged PAR, we also calculated the *OER* for fixed intensities of light (Figure 7E). We also calculated the standard parameters characterizing the photosynthesis-irradiance curves resulting from this model and tabulated them in Table 1. The parameter 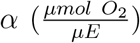 is the initial slope of the PI-curve,the parameter *P*_max_ (*μmol O*_2_ *m*^−2^*s*^−1^) is the maximum photosynthetic productivity determined by the peak position of the curves, the parameter *I_k_* (*μEm*^−2^ *s*^−1^) is the light level at which the system reaches half its maximum productivity, and the parameter 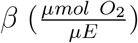 is the photoinhibition parameter and is the slope of the curve at high intensities.

**Table 1:** Photosynthesis-irradiance parameters calculated from radiative transfer model coupled to estimated quantum efficiency as a function of irradiance.

| | $\alpha \left( \frac{\mu\text{mol O}_2 \text{ m}^{-2} \text{ s}^{-1}}{\mu\text{E m}^{-2} \text{ s}^{-1}} \right)$ | $P_m \left( \mu\text{mol O}_2 \text{ m}^{-2} \text{ s}^{-1} \right)$ | $I_k \left( \mu\text{E m}^{-2} \text{ s}^{-1} \right)$ | $\beta \left( \frac{\mu\text{mol O}_2 \text{ m}^{-2} \text{ s}^{-1}}{\mu\text{E m}^{-2} \text{ s}^{-1}} \right)$ |
| --- | --- | --- | --- | --- |
| suspension | 4.5E-2 | 5.51 | 1.22E+02 | -1.98E-08 |
| small clam | 2.4E-2 | 302.10 | 1.27E+04 | -1.64E-05 |
| medium clam | 7.8E-2 | 1246.74 | 1.59E+04 | -7.28E-05 |
| large clam | 9.0E-2 | 1335.27 | 1.48E+04 | -6.50.6E-04 |

## Results

### Fluorescence transients

We measured fluorescence transients *in hospite* using a novel two-fiber optical microprobe coupled to a high-intensity 455 nm LED light source and PMT (Figure 4). As anticipated, we observed a log-linear onset of fluorescence (the “O” stage of the OJIP transient^1^); followed by a slight increase around 1 ms (the “J” stage); additional sample-dependent structure around 100 ms (the “I” stage); followed by a plateau around 1 ms (the “P” stage). An unanticipated complication of these experiments was that, as hypothesized, the microalgae embedded in the center of the giant clam system are indeed often very relaxed and highly quantum efficient. Our system performed extremely well, with high signal:noise in measuring the fluorescence transients of green leaves and zooxanthellae grown in culture flasks under moderate light levels (Figure 4A,B,C). In comparison, however, the fluorescence transients detected in the central regions of the whole clam tissue were of very low intensity, with correspondingly high signal:noise, and were also very slowly evolving (examples in Figure 4D).

### Tissue and cell suspension absorbance

In general, clam tissue exhibited high absorbance at all visible wavelengths, showing a maximum value around 0.9 at 400 nm and decreasing roughly linearly to a value of around 0.85 at 700 nm (Figure 5B). In comparison, a green leaf from a bamboo plant had a similar value at 400 nm, the characteristic local minimum in absorbance at around 550 nm of 0.85, and a minimum of 0.8 at 700 nm (Figure 5B). The serial dilution of a culture of suspended *Symbiodinium* measured in the same double-sphere geometry gave a single-cell absorbance cross-section of 56 μm^2^ at 460 nm (Figure 5C).

### Quantum efficiency

Using the fluorescence transient data measured in central regions of the tissue, we calculated the local quantum efficiency as a function of depth in the tissue. The quantum efficiencies of clam tissue were quite high. In general, we were surprised by the lack of dependence of the algal quantum efficiency on the vertical position in the tissue; the most shaded lower portions of the tissue were as efficient as centrally located cells, with some decrease in efficiency at the very top, most irradiated portion of the tissue. In the experiments where we succeeded in measuring three different vertical positions in the same sample, the measured efficiencies at the bottom and center positions of the tissue were very similar, ranging from from 0.55 to 0.89 with a median of 0.7 under actinic illumination (2000 μE μE m^−2^s^−1^), and ranged from 0.42 to 0.74 with a median of 0.6 under saturating illumination (5000 μE m^−2^s^−1^). Quantum efficiencies at the top of the tissue in this set of measurements were somewhat lower, ranging from 0.48 to 0.7 independent of illumination intensity (Figure 6A).

The majority of our experiments using this technique measured one vertical position per sample, due largely to constraints around the unexpectedly long time required to perform this experiment. Considering averages for single species under actinic illumination, *T. crocea* had a median value of 0.8, while under saturating illumination, this value decreased to 0.55. *T. derasa* showed a median efficiency under actinic illumination of 0.79 and a saturating value of 0.66; *T. gigas* had an actinic median of 0.8, and a saturating value of 0.52; the value for *H. hippopus* under actinic light was 0.87, while the saturating value was 0.72; *T. maxima* actinic value was 0.66, while the saturating value for this species was 0.52; and the *T. squamosa* actinic value was 0.6 while the saturating value was 0.45. In general, these values are high, and particularly so for organisms grown in the open at high tropical irradiance levels. Trends in the difference between actinic and saturating irradiance are clear and consistent. Caution should probably be used in interpreting any of the differences between species as significant given low sample numbers and unavoidable differences in the details of animal acquisition and handling. However, it is notable that the species found in shallower and more lensed habitats in Palau, for which we also have the most measurements, *T. crocea* and *T. derasa*, consistently had the highest measured efficiencies. In contrast, *T. maxima* was reliably lower, and is found in deeper, offshore wall habitats where lensing is slower and less intense. Relative to values reported in the literature, the decay in quantum efficiency with irradiance was surprisingly modest; the average for all measurements under the 2000 μE m^−2^s^−1^ incident light was 0.7, while at the 5000 μE m^−2^s^−1^ level it was 0.61 (Figure 6B). In general, these values are more consistent with a system grown and acclimated to low light intensities found in deep water, and not the extremely high intensities encountered on the shallow coral reefs where giant clams are found.

### Productivity models

Our goal was to model the productivity and efficiency of the clam system as if it were a material for energy transduction. We used two different light-exposure scenarios to estimate different performance parameters. In the first scenario, we calculated photosynthesis as a function of the irradiance at the system’s surface up to the extreme irradiances found in wave-lensed flashes. This allowed us to predict the system’s canonical photosynthesis-irradiance parameters P_max_, I_k_, *α* and *β* (Figure 7E, Table 1) at the very high irradiances at which a decay in efficiency ultimately occurred. In the second scenario, we predicted the areal, daily productivity of the system. In this scenario, the light increased smoothly from sunrise until noon and then decreased smoothly until sunset, but there were no pulses due to wave-lensing, and we integrated over 24 hours to find the system’s daily performance (Figure 7F).

Both productivity estimates use an established method^13^ to extrapolate the effective quantum efficiencies of a system using experimental photosynthesis-irradiance curves, in this case for whole clams.^4^ In Figure 7, the PI-curve data^4^ (thick lines) are plotted with the extrapolated quantum efficiency (dotted lines), and the resulting back-calculation to the underlying PI-curve data (thin lines). The medium- and large-size clams exhibited a more gradual increase in gross photosynthesis with irradiance, and the experimental data did not exhibit a clear maximum or asymptote value in oxygen evolution, even at the experimental irradiance of 3000 μE m^−2^s^−1^(three times greater than maximum time-averaged environmental downwelling irradiance) (Figure 7D). In the corresponding quantum efficiency relations, there are marked differences between the suspended zooxanthellae and the smallest clam, which have steep initial slopes, compared to the medium- and large-size clam systems, which show effective efficiencies of the system greater than 10% at irradiances of greater than 1000 μE m^−2^s^−1^at the system’s surface (Figure 7E). It is important to note that for any given surface irradiance, the local irradiances at single cells are on average around ten-fold less, given the process of radiative transfer through the system (Figure 7C). The performance of the system is, then, the sum of single-cell behavior in “diluted” light within the depth of the tissue. It is the specifics of radiative transfer through the system, with light initially scattered at small, forward angles by the surficial iridocytes and then interacting with the vertical faces of the algal pillars, that leads to the apparent, extremely high predicted effective efficiencies at extremely high surface irradiances (Figure 7E). As proof-of-concept for this approach, these calculations produces similar areal oxygen evolution for our suspended algal sample as for a soybean leaf under saturating irradiance (5.51 *μ* mol O_2_ m^−2^ s^−1^ vs. 10 *μ* mol O_2_ m^−2^ s^−1^).^20^ Surprisingly, large clams reach maximum oxygen evolution rate of 1335 *μ*mol O_2_ m^−2^ s^−1^ at an irradiance of 212,000 μE m^−2^s^−1^, with a value of *I_k_* of 148,000 μE m^−2^s^−1^(Figure 7E). The same parameters for suspended algae, and the small and medium clam were correspondingly and systematically lower, and are reported in Table 1.

In the second model incorporating a circadian cycle of environmental irradiance, the three clam models were five to twelve times more productive compared to a system with the same number and average density of algae organized randomly. As the clam tissue thickness increased from 0.3 to 4.5 mm, the areal productivity increased from 1.5 to 3.5 moles O_2_ m^−2^ day^−1^ (Figure 7F). However, O_2_ evolution per cell decreased with tissue thickness from about 1990 to 790 pmol O_2_ cell^−1^ hr^−1^ (Figure 7F). When compared to a physical optimum, where all cells in the model converted all absorbed light at 100% quantum efficiency, the medium clam reached 93% of the physical optimum, and the large clam reached 88% of the physical optimum on a per area basis, and 94% and 92% of the optimum on a per-cell basis, respectively. In contrast, a random layer of with the same number of cells and average density reached 6% of the physical optimum on both a per-area and a per-cell basis (Figure 7F). On a per-cell basis, the small clam reaches 84% of the physical optimum, and the large clam reaches 92% of the optimum (Figure 7F).

With the combined effects of high per-cell efficiencies and low homogeneous light levels, the clam models (small, medium, and large) were 5-12 times more productive on a per-area basis than a system with homogeneous algae distributions. The clams models increased in areal productivity as the tissue thickness increased from 1.5 to about 3.5 moles O_2_ m^−2^ day^−1^. The tradeoff for increasing the tissue thickness is that both the optimum and the observed per-cell productivity decreases.

## Discussion

We were surprised by the high quantum efficiencies we measured in clams raised in the extreme irradiances of shallow Palauan reefs. Experimental quantum efficiencies were consistently very high, even at stimulation intensities much higher than average downwelling PAR (even our “actinic” stimulation was a higher-than-natural 2000 μE m^−2^ s^−1^, but necessary to generate enough fluorescent photons to measure in most experiments). In particular, *T. crocea* had an average efficiency of 80% at an irradiance of 2000 μE m^−2^s^−1^and in some experiments it was as high as 90%. Efficiency values were higher in *T. crocea*,*T. derasa*, *T. gigas*, and *H. hippopus* than in *T. maxima* or *T. squamosa*. The *T. crocea*, *T. gigas*, and *T. derasa* we worked with in Palau characteristically grew in super-tidal coral heads, staghorn coral beds, and seagrass areas, respectively. All three of these habitats are subject to intense wave-lensing patterns on clear days. In contrast, *T. maxima* and *T. squamosa* tend to be found deeper, in clear, relatively off-shore waters of fore-reefs and vertical walls, where irradiances can be high but wave-lensing is attenuated.

We interpret these experimental fluorescence transient data to be good proof of principle that the photosystems within mature, wild giant clams from shallow reefs can be extremely relaxed and absorbing throughout the depth of the clam tissue. Our experimentally measured quantum efficiencies actually far exceed the efficiencies incorporated in the productivity model in the interior of the clam tssue. However, because our experimental data are also variable between species and within the tissue, it was difficult assign efficiency values as a function of depth within the tissue. Therefore, we used efficiencies extrapolated from whole-system experimental photosynthesis-irradiance curves coupled to a model of irradiance as a function of depth within the tissue.

This approach allowed us to estimate photosynthetic productivity over a wide range of irradiances, finding standard parameters like assimilation number,*P_m_*, the initial slope of P-I, *α*, the photoadaptation parameter, *I_k_*, and the photoinhibition parameter, *β*,.^21^ We could also then meaningfully scale productivity by the surface area and cell density of the system, the important parameters for any engineered devices using similar physical principles to the clam.

These values are reported in Table 1. Surprisingly, the photoadaptation parameter *I_k_*, or the light level at which the system reaches half its maximum productivity, is greater than the solar constant for all the clams. PAR at the top of the atmosphere at the equator is about 3100 μE m^−2^s^−1^, while *I_k_* for all the clam models was greater than 12,000 μE m^−2^s^−1^.

The maximum rate of oxygen evolution per tissue area, or the area-normalized assimilation number, *P_m_*, of 1335 *μ*mol O_2_ m^−2^s^−1^occurred at an irradiance of about 1.5×10 ^4^ μE m^−2^s^−1^, or about fifteen times brighter than average noontime PAR in Palau of 1000 μE m^−2^s^−1^(Figure 7E). We initially assumed we had made a mistake in our calculations. This finding suggests that clams are adapted to efficiently photoconvert irradiances much higher than the average irradiance of their habitat, and higher than the solar constant, or the flux at the top of the atmosphere at the equator (about 3100 μE m^−2^s^−1^). Experimental wave-lensing irradiance data from Santa Barbara Channel show pulses in downwelling irradiance ten times greater than the average background lasting several milliseconds and occurring a few times a minute.^22^ This finding sets a lower bound for expectations around intense flashes in the clam’s habitat, where we expect extreme pulses to be even brighter and more frequent under the clear skies and breezy conditions that are common there. Indeed, the largest clam’s productivity starts to level off at an irradiance around 15 times the short-time-average noontime environmental irradiance. This observation suggests that this clam is primarily adapted to flashes that are 15 times brighter than the background irradiance, an intensity well within the expectations of the existing wave-lensing models.

This apparent adaptation to the extreme radiances in wave-lensed pulses peaks means that the clams can be highly efficient at biological photoconversion at intensities much higher than the solar constant. Remarkably, the clam tissue is likely close to physically optimal for a biological system. When we compared the performance of the clam tissue to one where cells can convert light energy to chemical energy at a quantum efficiency of unity, the clam system came within 15% of those values. In contrast, the system with the same number of cells arranged randomly was less than 10% of this optimum value on either a per-cell basis or a whole-system basis (Figure 7F). The physical principles realized by giant clams may in turn then be important for understanding how to engineer efficiency and robustness to photodamage into both bio- and organic based solar cell technologies under solar concentration. The clam system plausibly could come within 10% of perfect efficiency of phototransduction even at intensities several times greater than the average downwelling intensity at the equator. Presumably the physical limit of the clams’ strategy is when the per-cell irradiance in the thickest systems falls below the photosynthesis-respiration limit for a single cell, or the light compensation point of the clam system in analogy to the productivity of the open ocean. However, even the largest clams had several hundred-fold greater per-cell productivity than the system with homogeneous organization, suggesting that clams exist far from the light compensation point of the resident zooxanthellae and ever greater areal productivities are in principle possible in engineered systems.

## Acknowledgement

We thank the staff and scientists of Palau International Coral Reef Center, in particular Ikelau Otto, for support working with giant clams; Dr. Yim Golbuu for mentorship of LFR; aquarist Asap Bukurrou for his wild clam expertise; and clam farmer Kenneth Mereb for helping us with clam procurement. This work was supported by NSF INSPIRE award 1343159 and a David and Lucile Packard Foundation fellowship to AMS.

## Supporting Information Available

Data and matlab codes used to analyze and create the figures from this manuscript are available on Dryad: https://doi.org/10.5061/dryad.xsj3tx9k4

## Supplemental Detailed Methods

### Double optical-microprobe construction

In order to measure photosynthetic transients within the highly absorbing living tissue, we modified an optical microprobe technique to incorporate two optical fibers. One of these fibers stimulates the tissue with 455 nm light, and the second fiber receives the resulting time-resolved chlorophyll fluorescence response. We fabricated these probes as follows. The termination of two ends of 100-μm silica core fiber optic assemblies (Ocean Optics, Dunedin, FL, USA) were removed. The end of each fiber was tapered using the following method: First, the fiber jacketing was stripped away and the fiber’s polyimide buffer was removed for a distance of 5 cm from the fiber’s end using a butane torch. Then, a 10 g weight was attached to the end of the fiber and the fiber was then pulled, narrowing the diameter, upon heating with a butane torch. Then, the narrowed region of the fiber was then cut using carborundum paper, to yield a flat fiber end with a diameter of 30–50 μm. Finally, the sides of the narrowed fiber were painted with a film opaquing pen to prevent stray light from entering, while leaving a small transparent opening at the fiber’s tip. The end diameter of each tapered fiber was characterized using a microscope. The absolute intensity of light output from each tapered fiber was measured using an integrating sphere (AvaSphere-50-LS-HAL-CAL) and an Avantes USB spectrometer (AvaSpec ULS2048L-USB2) calibrated to a NIST-traceable halogen lamp source (AvaLight-HAL-CAL-Mini). The ratio of the diameter of the tapered optical fiber to the integrating sphere port can then be used to calculate intensity of the excitation light source in units of PAR, or μE m^−2^s^−1^. The two tapered and optically characterized fibers were then carefully aligned next to each other in the tip of a pulled glass Pasteur pipette and secured using a drop of cyanoacrylate glue, leaving only 2–3 mm of bare optical fibers protruding Figure 3.

### Tissue preparation and fluorescence transient measurement

We measured fluorescence transients within the living tissue of several species of giant clam. *Tridacna derasa*, *T. maxima*, *T. crocea*, *Hippopus hippopus*, *T. squamosa* and *T. gigas* were purchased from clam aquaculture facilities in Palau and kept alive in running seawater at the Palau International Coral Reef Center (PICRC) in Koror, Palau. To prepare clam samples for optical measurements, the clam shell adductor muscle was severed from one valve and the photosynthetic region of the mantle was trimmed away. We used a biopsy punch to obtain an 8 mm-diameter biopsy of the complete thickness of the mantle margin, which was 2–4 mm thick, depending on the animal. This circular piece of tissue was then placed in an 8-mm tape-covered hole in the bottom of a petri dish, and in-filled with room-temperature molten gelatin. This preparation was then refrigerated briefly until the gelatin solidified. The resulting rigidity of the gelatin provided a transparent matrix that was sufficiently rigid to immobilize the clam tissue in the dish during the experiment. The Petri dish containing the clam sample was then mounted on a stage attached to a micromanipulator and positioned to center the hole in the bottom of the dish around the fiber optic probe tip, which was mounted vertically and held stationary throughout the experiment. We then lowered the dish in fixed increments onto the double-optical probe, pushing the probe through the clam tissue (Figure 3A).

Stimulating 455 nm light was provided by a ThorLabs M455F1 fiber optic-coupled high power LED coupled to one of the two fibers in the probe. The fluorescence transient was measured in turn using the second fiber from the probe connected to the SMA port of a ThorLabs photomutliplier tube (PMTSS). The photomultiplier tube (PMT) signal was detected using an GWINSTEK oscilloscope (GDS-2072E). The stimulating blue light was short-pass filtered through an Edmond Optics 600 nm OD4 short-pass filter before reaching the sample, and the fluorescent response was long-pass filtered using an Edmond Optics 650 nm OD4 long-pass filter before reaching the PMT (Figure 3A). We measured fluorescence transients at different light stimulation intensities of 2000, 3500, and 5000 μE m^−2^s^−1^at the immediate surface of the probe. These stimulating intensities are higher than typically used in this type of experiment, however, in practice these intensities were required to generate measurable fluorescence responses specifically when measuring zooxanthellae within clams. We characterized the flux from each new individual probe using an AvaSphere-50-LS-HAL-CAL integrating sphere. For terminological convenience, we considered 2000 μE m^−2^s^−1^at the probe tip to be “actinic” and 5000 μE m^−2^s^−1^at the probe tip to be “saturating” in our subsequent analysis of these data, while also noting that these terms are typically defined at much lower intensities.

The dynamic portion of the chlorophyll fluorescence transient occurs over timescales ranging from a few microseconds to a few hundred milliseconds, such that measurement apparatus must similarly operate on microsecond time scales. We determined the rise time of the M455F1 LED light source, when triggered using ThorLabs DC2100 controller, to be 5 μs when the system was run in pulse-width modulation mode. We then programmed the intensity (via current) and frequency of the LED using the manufacturer’s software.

We maximized the signal from the photomultiplier tube while keeping the usable measurement response time under 20 μs by terminating the PMT with a 100 kΩ resistor. The fluorescence signal from the photomultiplier tube was recorded with the oscilloscope at a time resolution of 2 μs per recorded time point. The oscilloscope’s recording was triggered manually immediately before the excitation LED was switched on, and the resulting fluorescence signal was recorded on the oscilloscope for 2 s.

Prior to each measurement, the clam tissue sample was dark adapted at room temperature under a box for 15 minutes. Due to the sensitivity of the PMT and to ensure all signal was from chlorophyll fluorescence, all measurements were performed in a dark room.

### Fluorescence transient data analysis

To calculate the maximum quantum yield of zooxanthellae *in hospite* resulting from these measurements of the fluorescence transient, we started by identifying the zero-time point of each measurement where signal started to rise by eye. Then, using purpose-written Matlab code, we used a median filter to smooth the rapidly sampled oscilloscope data while preserving the time-dependence of the fluorescence signal. The quantum efficiency of the algae near the optical probe is characterized by the minimum chlorophyll fluorescence at the zero-time point (*F*_0_) and the maximum fluorescence at a time greater than 1 s (*F_M_*). To find *F*_0_ from the fluorescence transient data, we made a linear fit to the fluorescence signal between 40 and 120 μs and calculated its intercept at the zero-time point.^14^ *F_M_* is then the steady-state fluorescence value after 1 s when the signal was averaged over 5 ms intervals. The maximum quantum yield, *q* is then calculated using the definition 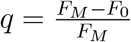 ^1^ (Figure 4).

### Monte-Carlo Modeling

The Monte Carlo algorithm used in this paper is described in detail by Holt et al.^**?**^ Briefly, we generated three-dimensional coordinates of algae and iridocytes based on two-dimensional panoramic light micrographs and dark field light microscope images. These coordinates were used in a Monte Carlo algorithm of light propagation where each cell type scattered photon packets according to modeled phase functions based either on Mie scattering (algae) or Discrete Dipole Analysis (iridocytes).^2,15^ Trajectories of 90,000 photon packets were modeled in single runs, resulting in 1%–5% uncertainty around single-cell absorption values. Using this Monte Carlo model, we calculated the average per-cell absorption as a function of tissue depth for three scenarios, keeping all geometric parameters the same except for the maximum depth of the tissue. We generated three-dimensional cell coordinates for algal pillars of about 4.6 mm depth, consistent with large individual clams (1000’s of grams wet weight). We then truncated these coordinates twice, once at 1.2 mm (medium-sized clam with 18.2 g wet weight) and again at 0.350 mm (small-sized clam with a 0.014 g wet weight), such that the positions of algae and iridocyte cells were equivalent for each model up until the maximum tissue depth. This set of models is consistent with our observations of clams of different size- and age-classes, where the two-dimensional arrangement of the pillars within the mantle tissue doesn’t change as clams grow, but the mantle tissue becomes thicker with shell length. It is also consistent with the scaling relations between shell length, wet weight and zooxanthellae cell number cited in Fisher et al.^4^ The calculated whole-tissue abosrbance of these model systems is also in agreement with experimental measurements of whole-tissue absorbance measured by doubled integrating spheres, described below.

### Clam tissue and single-cell absorbance measurements

To experimentally measure the total amount of light absorbed in the clam mantle tissue, we used a doubled integrating sphere approach. We prepared mantle biopsies embedded in gelatin as described above, and sandwiched the slab of gelatin containing the clam tissue between two high-precision windows with antireflection coating. This preparation was then mounted in a ThorLabs translation mount (ST1XY-S). The translation mount with the piece of tissue was pressure mounted between two 2-inch ThorLabs integrating spheres (IS200-4). The light source for this experiment was a ThorLabs fiber-optic coupled white LED (MCWHF2) secured to one integrating sphere using cage mounts. Light measured from the sphere with the light source mounted is then primarily back-scattered from the sample, and light measured from the sphere without the light source is light primarily transmitted by the sample. Fibers were mounted perpendicular to the light path on both integrating spheres to characterize this back-scattered and forward-scattered light. We centered the clam tissue in the spheres’ sample apertures using the translation mount and verified by eye that the tissue filled the ports of the integrating spheres such that all light incident on the clam tissue must pass through the tissue in order to reach the second integrating sphere. We standardized this measurement of clam tissue by characterizing the light in both spheres without a clam sample present. The signal missing in the second sphere when the clam tissue is present, relative to the signal when it is absent, is then the light that is absorbed by the tissue. We also characterized the absorptance of several green leaves in this manner (Figure 5A,B).

We also estimated the absorptance and transmittance of single zooxanthellae cells from a dilute suspension of cells. Cells grown in dilute culture were washed and serially diluted with growth medium. Cell suspensions were then placed in a 1 cm-pathlength quartz cuvette mounted between two integrating spheres (Figure 5A). The total reflectance and transmittance of the algae in solution was measured from the signal in the two spheres and normalized to a quartz cuvette filled with clean growth medium. The cell number density of each dilution was measured using a hemocytometer. We calculated the single-cell absorption cross-section from the least-squares solution to the absorbance (log of transmittance values at 460 nm) versus concentration data points (Figure 5C).

### Primary productivity model

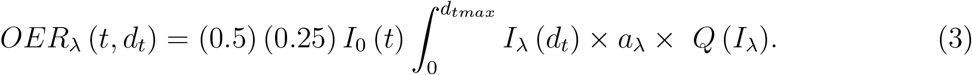

In this calculation, we extrapolated cellular quantum efficiency from photosynthesis-irradiance (PI) relationships of the clam system reported in the literature using the method of Eilers and Peeters.^4,13,17,18^ Using the nonlinear least-squares method as incorporated in Matlab, we fit experimental PI curves from the literature from a range of different sized, living Tridacnid clams and also from zooxanthellae isolated from this tissue to the equation

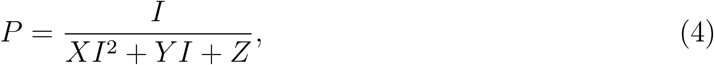

where *P* is the photosynthetic productivity, *I* is the irradiance, and *X*, *Y*, and *Z* are fit parameters. Using this fit, the relation to the photosynthetic quantum efficiency using the method of Yang et al.^18^ is:

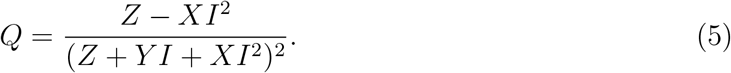

We then normalized the efficiency curves to maximum efficiency at zero irradiance.

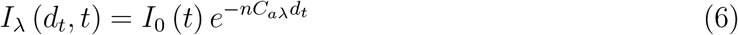

We obtained the time-dependent environmental PAR (*I*_0_ (*t*)) from a database from the Pacific Islands Ocean Observing System^19^ which tracked PAR in Koror, Palau for many months throughout the year 2012. We averaged PAR as a function of time of day over 30 days in August to get an estimate of daily PAR for each hour over a 24-hour period. PAR covers wavelengths from 400-700 nm but the rest of our calculations and measurements were based on the same amount of total energy propagating through the system at a single wavelength at 450 nm. This simplification only influences scattering from iridocytes and between cells since our model does not incorporate fluorescence or secondary effects of wavelength on photosystem efficiencies. We calculated *OER*_*λ*(*t*)_ over a 24-hour period using *Q* (*I_λ_*) for the four systems mentioned above.

To model the effects of intense pulses from wavelensing and the performance of the system at intensities higher than short-time averaged PAR, we also calculated the *OER* for fixed intensities of light (Figure 7E). We also calculated the standard parameters characterizing the photosynthesis-irradiance curves resulting from this model and tabulated them in Table 1. The parameter 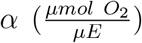 is the initial slope of the PI-curve and was determined by fitting a line to the data between irradiances of 0 to 100 *μE m*^−2^ *s*^−1^. The parameter P_max_ (*μmol O*_2_ *m*^−2^*s*^−1^) is the maximum photosynthetic productivity determined by the peak position of the curves. The parameter *I_k_* (*μEm*^−2^ *s*^−1^) is the light level at which the system reaches half its maximum productivity and is calculated by dividing *P*_max_ by *α*. The parameter 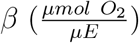 is the photoinhibition parameter and is the slope of the curve at high intensities. We calculated *β* by fitting a line to the data between irradiances of 280,000 - 1,000,000 *μEm*^−2^*s*^−1^.

No unexpected or unusually high safety hazards were encountered during the course of these experiments.

## Notes

### Competing Interest Statement

The authors have declared no competing interest.

